# DIA-BERT: pre-trained end-to-end transformer models for enhanced DIA proteomics data analysis

**DOI:** 10.1101/2024.11.11.622930

**Authors:** Zhiwei Liu, Pu Liu, Yingying Sun, Zongxiang Nie, Xiaofan Zhang, Yuqi Zhang, Yi Chen, Tiannan Guo

## Abstract

Data-independent acquisition mass spectrometry (DIA-MS) plays an increasingly important role in quantitative proteomics. Here, we introduce DIA-BERT, a software tool that leverages a transformer-based pre-trained artificial intelligence (AI) model for the analysis of DIA proteomics data. Over 276 million of high-quality peptide precursors extracted from existing DIA-MS files were used for training the identification model, while 34 million peptide precursors from synthetic DIA-MS files were used for training the quantification model. Compared to DIA-NN, DIA-BERT led to on average 54% more protein identifications and 37% more peptide precursors in five different human cancer (cervical cancer, pancreatic adenocarcinoma, myosarcoma, gallbladder cancer, and gastric carcinoma) sample sets with a high degree of quantitative accuracy. This study highlights the potential of utilizing pre-trained models and synthetic datasets to advance DIA proteomics analysis.

## Main

Analyzing data-independent acquisition mass spectrometry (DIA-MS) based proteomics data remains a formidable challenge^1-3^. Pioneering DIA identification tools, such as OpenSWATH^4^ and Spectronaut^5^, employ a targeted data analysis strategy similar to that used in multiple reaction monitoring (MRM), like mProphet^6^ and Skyline^7^. In this strategy, peak groups, instead of mass spectra, are extracted for scoring, significantly reducing data complexity, and largely preserving the major ion trace and retention information of peptide precursors and top N fragment ions. Recent DIA identification tools, such as DIA-NN^8^, MaxDIA^9^, Specter^10^ and DreamDIA^11^, typically employ a multi-stage approach. They initially score peak groups with relatively simple rules or models and then refine the selection through an iterative procedure. This process leverages an expanding array of features, progressively enhancing the discriminative power of the scoring model to improve selection accuracy. Nevertheless, current DIA identification tools face limitations. They train independent scoring models for each mass spectrometry file, and due to the limited training sample size from each file, these tools typically rely on conventional machine learning models with limited capacity to avoid overfitting. Training the scoring model separately for each MS file prevents it from leveraging information from other files. As a result, they often fail to adequately capture intricate peptide-peak-group matching patterns. Their reliance on manually engineered features such correlations between different metrics in current methodologies further limits the overall analytical performance. Additionally, training the representation model and the scoring model independently, or relying on handcrafted features, prevents an information feedback loop between these two stages, thus missing opportunities for joint learning.

Emerging approaches, such as AlphaTri^12^and Alpha-XIC^13^, attempt to use more complex deep learning models specifically designed for MS spectra to capture intricate peptide-peak-group matching patterns. However, their scoring model is still trained independently for each individual mass spectrometry file, which limits the complexity of the models. It uses the relatively simple two-layer Long Short-Term Memory (LSTM)^14^ models with limited scoring performance. DreamDIA^11^ advances this exploration further. It first trains a deep representation model using training samples extracted from multiple mass spectrometry files. The learned representation is then fed into an XGBoost^15^-based scoring model, along with some manual features, to filter peak groups. While the representation model is trained on one million representative spectral matrices (RSMs), the shallow XGBoost scoring model is still trained separately for each mass spectrometry file and the representation model and the scoring model are still trained independently, limiting its power.

BERT^16^, an acronym for Bidirectional Encoder Representations from Transformers, stands at the forefront of natural language processing (NLP) breakthroughs. Here, we introduce DIA-BERT (**Figure 1a, Methods**), a transformer-based pre-trained AI model tailored for analyzing DIA-MS proteomics data. We present a DIA identification algorithm leveraging an encoder-only transformer model. The models in the initial (pre-filtering model) and subsequent (pre-trained scoring model) stages undergo training on extensive datasets comprising 276 million and 318 million training instances sourced from 952 DIA proteomics files from various human specimens, respectively. The identification model was trained using peak groups extracted from the 952 DIA files by DIA-NN^8^ and Spectronaut^5^ using library-based strategy. The breadth of these datasets equips our models with exceptional generalization prowess. Harnessing the robust data modeling capabilities inherent in convolutional and transformer networks empowers our models to adeptly capture intricate peptide-peak-group matching patterns. Our model undergoes end-to-end training, eliminating the necessity for a separate handcrafted feature extraction or representation learning step required by solutions such as DreamDIA^11^ and other conventional machine learning-based methods. The identification model is pre-trained on a large dataset and then fine-tuned on each mass spectrometry file. Pre-training minimizes the problem of overfitting due to insufficient samples that can arise from the independent training of each file. Fine-tuning enables the model to capture the unique characteristics of each file. This unified training process enables our model to concurrently optimize representation learning and scoring model learning, leading to enhanced identification sensitivity.

**Figure 1.**
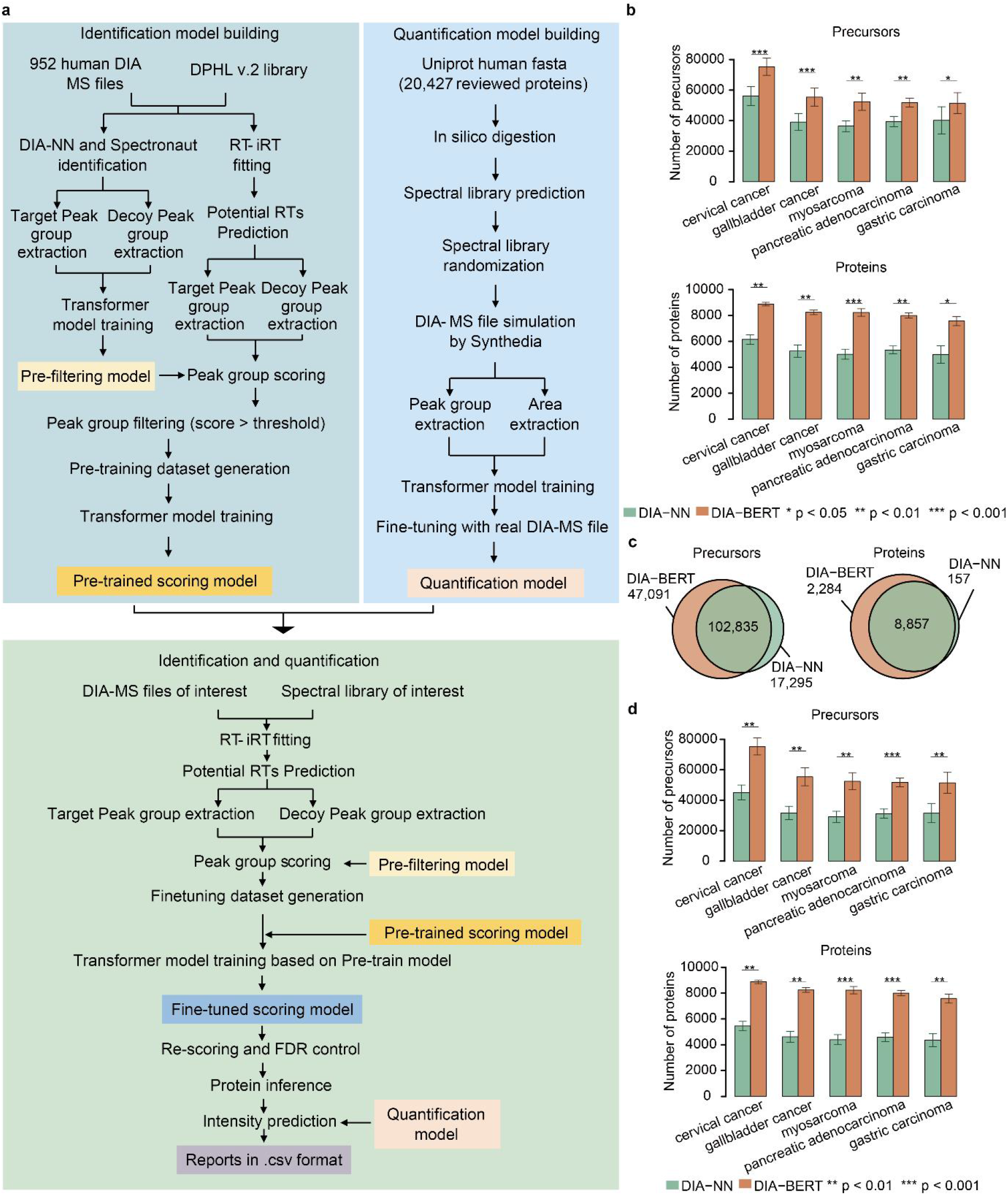
DIA-BERT workflow and its performance on different human cancer data set. **a**, The workflow of DIA-BERT. **b**, Comparison of peptide and protein identifications between DIA-BERT and DIA-NN (library-based mode) in human proteome data set. **c**, The overlap of identified human tissue peptide precursors or proteins using DIA-BERT and DIA-NN (library-based mode). **d**, Comparison of peptide and protein identifications between DIA-BERT and DIA-NN (library-free mode) in human proteome data set.

To assess the performance of DIA-BERT, we first used a DIA proteome dataset of various human tumor tissues^17^, comprising three samples each from five different cancers (cervical cancer, pancreatic adenocarcinoma, myosarcoma, gallbladder cancer, and gastric carcinoma), sourced from the iProx database (IPX0001981000). The DIA pan-human library (DPHL) v.2 was employed as the spectral library^18^. DIA-NN is the currently best performing DIA proteome software in multiple evaluation studies^19, 20^, therefore, we compared our software tool with DIA-NN. As shown in **Figure 1b** and **Table S1**, DIA-BERT led to significantly more peptide precursors and protein identifications compared to DIA-NN in all DIA files. On average, DIA-BERT identified 37% more peptide precursors and 54% more proteins than DIA-NN. The number of peptide precursors or proteins identified by DIA-BERT is significantly (two-sided paired Student’s *t* test, p-value < 0.05) higher than those identified by DIA-NN in five different human cancer sample sets (**Figure 1b**). DIA-BERT re-called 86% of peptide precursors and 98% of proteins identified by DIA-NN (**Figure 1c, Table S2**). We evaluated the identification consistency across samples in five different human cancer sample sets. The results showed that DIA-BERT consistently identified 62% of precursors and 86% of proteins in 2/3 or more of the samples on average, compared to 61% and 79% for DIA-NN version 1.9 in library-based mode (**Table S2**). We also assessed the performance of DIA-BERT against DIA-NN in library-free mode. Compared to DIA-NN in library-free mode, DIA-BERT identified 76% more proteins and 71% more peptide precursors on average using the DIA files from five human tumors (**Figure 1d, Table S1**).

We further dived into the unique peptide precursors and proteins identified by DIA-BERT compared to DIA-NN. The frequency of the number of peptide precursors per protein between DIA-BERT and DIA-NN showed that DIA-BERT can identify 24% more proteins with fewer than 10 peptide precursors, and 24% more proteins with at 10 or more peptide precursors (**Figure S1a**). We hypothesized that DIA-BERT has the ability to identify more low abundant proteins. To test this hypothesis, we compared the abundance of unique and common peptide precursors/proteins. The abundance of unique precursors/proteins was significantly (two-sided unpaired Student’s *t* test, p-value < 0.05) lower than that of common peptide precursors and proteins (**Figure S1b**). This phenomenon is consistent with DIA-NN (**Figure S1c**), indicating that DIA-BERT has an improved ability to identify low-abundance peptide precursors and proteins while maintaining a similar ability to identify high-abundance peptide precursors and proteins.

We further evaluated the performance of DIA-BERT in a three-species dataset^19, 21^ (human, yeast, and *C. elegans*) sourced from the PRIDE database (PXD016647). DIA-BERT achieved a 9% enhancement in protein identification and a 19% improvement in peptide precursor coverage compared to DIA-NN (**Figure S2a, Table S1**). The number of identified peptide precursors and proteins by DIA-BERT significantly exceeded that of DIA-NN, as indicated by a two-sided paired Student’s *t* test (p-value < 0.05) (**Figure S2a**). DIA-BERT re-called over 94% of peptide precursors and 94% of proteins identified by DIA-NN (**Figure S2b**). Compared to DIA-NN, the abundance of peptide precursors and proteins exclusively identified by DIA-BERT was significantly lower than that of peptide precursors and proteins identified in both DIA-BERT and DIA-NN (two-sided unpaired Student’s *t* test, p-value < 0.05) (**Figure S2c**). This mirrors observations in DIA-NN (**Figure S2d**), suggesting DIA-BERT’s enhanced capability in identifying low-abundance peptide precursor/proteins in library-based mode. This strength extends not only to human proteomes but also to proteomes of three species. Compared to human tumor tissue datasets (DPHL v.2 library size: 601,982 precursors), the improvement in identification ability for the three-species dataset is less pronounced. This may be attributed to the smaller library size (262,227 precursors), which might have not reached saturation. DIA-BERT has already identified 58.2%, 73.3%, 55.9% precursors in the library for human, yeast, *C. elegans* respectively. We also assessed the performance of DIA-BERT against DIA-NN in library-free mode. Compared to DIA-NN in library-free mode, DIA-BERT identified 34% more proteins and 57% more peptide precursors on average using the DIA files from three-species DIA data set (**Table S1**).

Next, to ensure precise quantification of peptide precursors and proteins, we developed a novel peak area estimation algorithm based on transformer models (see **Methods**). Current DIA data analysis methods rely on simplistic heuristic rules to compute peak areas^9^, followed by basic adjustments based on metrics like correlation coefficients between peaks within the same group. However, due to substantial noise interference in mass spectrometry signals and intricate mutual interference among multiple peak groups, these methods often yield unsatisfactory results. While downstream corrections can be applied using techniques such as MaxLFQ^22^ and DirectLFQ^23^ for cross-run analysis, the adverse effects of initial estimation errors remain challenging to fully mitigate.

Here, we used Synthedia^24^, a software package designed for simulating DIA data, to generate training DIA data annotated with TRUE area values. This data was then used to train a transformer-based peak area estimation model, aiming to enhance the accuracy of area estimation. The structure of our peak area estimation transformer model mirrors that of the model used for peak group scoring, with the key difference being the application of a regression loss function to estimate the peak area. This model processes the entire peak group’s information, utilizing data from all peaks within the group to mitigate the negative impact of noise interference on the current peak. The generated dataset includes numerous mutually interfering peak groups and random noise, enabling our model to effectively handle complex scenarios encountered in experimental mass spectrometry data. We modified the codes of Synthedia to accommodate various situations encountered in real MS data. We utilized a range of chromatographic and mass spectrometry parameters (see **Table S3**) to generate sufficient training data. In total, 360 mzML files (6.87 TB, comprising 34 million training instances) were simulated. The quantification model was finally fine-tuned with 36 existing DIA-MS files of human cell line samples.

To assess the quantification model, we utilized again the three-species data set^19, 21^ comprising human, yeast, and *C. elegans* samples with pairwise ratios of 1:1, 2:1, and 1:1.3, respectively. For single-file searches, DIA-BERT demonstrates quantification performance comparable to DIA-NN, with average Pearson correlation coefficients of 0.92 for peptide precursor quantification and 0.84 for protein quantification. For combined search, cross-run normalization was performed (see **Methods**). The median of log2-ratios predicted by DIA-BERT for peptide precursors (0.049, 0.84, and -0.36) and proteins (-0.05, 0.84, and -0.36) aligns consistently with the theoretical values (0, 1, and -0.38) for each species (**Figure 2**). DIA-BERT showed comparable median coefficient of variation (CV) values for peptides and proteins (4% and 8%, respectively, compared to 10% and 5% for DIA-NN).

**Figure 2.**
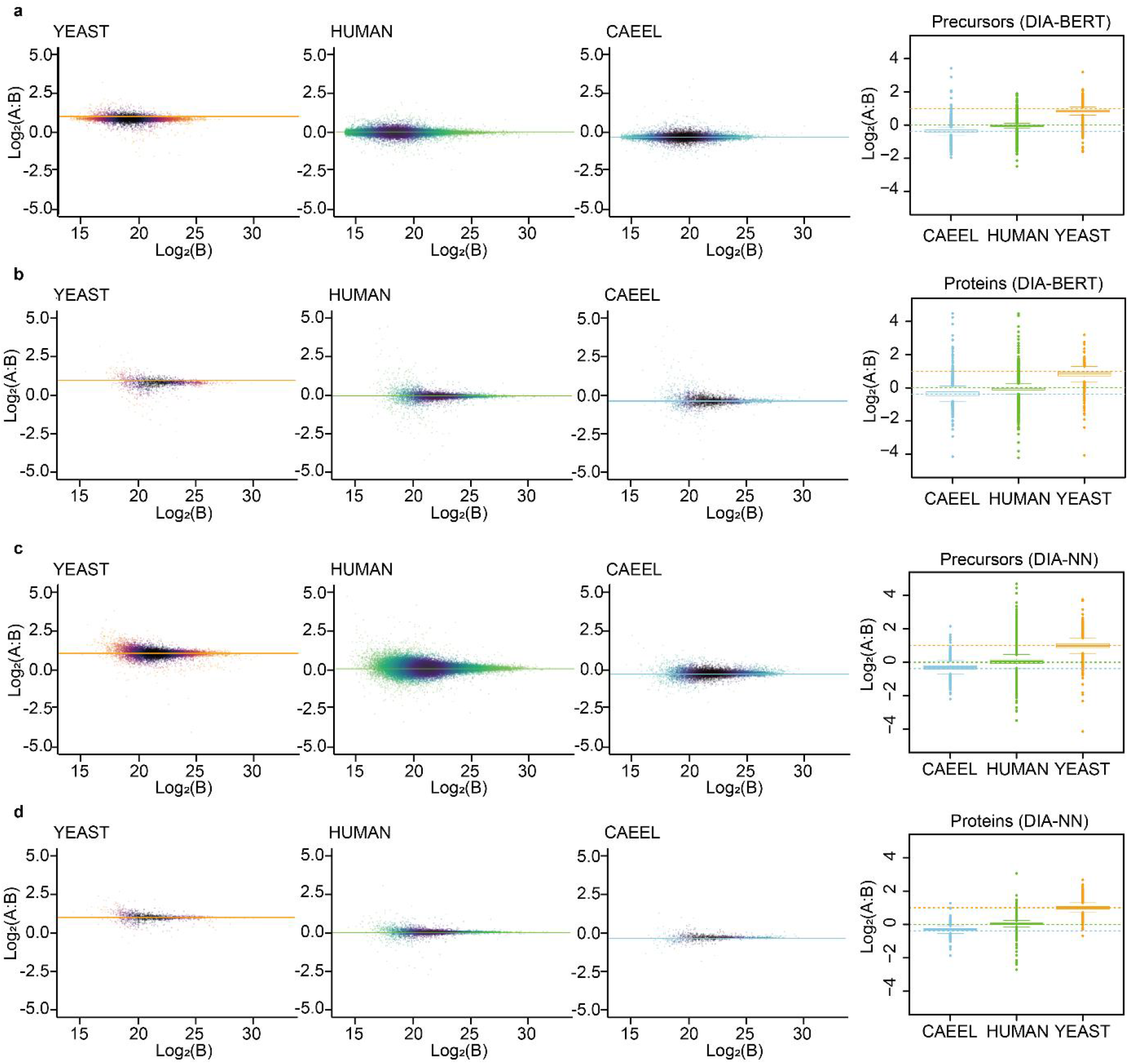
The quantification for three species in three species proteome data set (left three panel) and boxplot (right panel) at peptide precursor level (**a**) and protein level (**b**) using DIA-BERT, and peptide precursor level (**c**) and protein level (**d**) using DIA-NN. Orange color represents yeast; Green color represents human; Sky-blue color represents *C. elegans*.

DIA-BERT is user-friendly, featuring an intuitive graphical interface and providing results in CSV format. The graphical user interface (GUI) of DIA-BERT comprises four main sections: Input, Configuration, Output, Run Progress, and Log (**Figure S3**). These sections collectively streamline the analysis procedure. The Input section facilitates easy setup by allowing users to specify paths for raw files and customize the output directory. The Configuration section enables adjustments for settings such as the spectral library, run threads, decoy method, RT normalization, m/z unit, RT peak size, RT unit, instrument, batch size, batch score size, RT width, fine-tune score, cross-run quantification, and the number of epochs for model optimization. The Run Progress section monitors the analysis, allowing users to toggle between run and pause states while providing real-time progress updates. Finally, the Log section serves as a comprehensive runtime log, capturing essential events, state transitions, errors, warnings, and critical runtime details. This feature offers users valuable insights into the analysis process, aiding in comprehension and assessment.

In summary, the pre-trained transformer-based model introduced in DIA-BERT significantly improves identification of peptide precursors and proteins in DIA proteome data sets across different species samples. This study demonstrates that a pre-trained transformer-based model can enhance the analysis of DIA proteomics, offering greater potential for advancing understudied areas in proteomics. In principle, it is also feasible to extend DIA-BERT for the analysis of DIA data of metabolites and lipids. Additionally, we also established a transformer-based model for precise quantification of DIA data using simulated DIA datasets for training, paving the way for low-cost, large-scale training in proteomics without the need for complex and time-consuming experiments. Together, this study introduces a robust new software tool for DIA proteomic identification and quantification, highlighting the substantial potential of pre-trained models and simulated datasets in advancing DIA proteomics analysis.

This study has several limitations. Firstly, the utilization of DIA-BERT requires GPU computers, similar to other large models. Based on our experiments using NVIDIA A100 or V100 GPUs, a single file analysis generally takes around two hours to complete. Future software engineering enhancements are necessary to optimize resource utilization. Secondly, the identification model predominantly relies on training data from 952 DIA files obtained from human specimens. However, there is room for improvement by incorporating more DIA files from diverse species and mass spectrometry instruments. It is important to note that such enhancements will necessitate additional computational resources.

## Materials and Methods

### Mass spectrometry files used in training model

A total of 952 mass spectrometry files were collected, including 600 publicly available files from PRIDE and 352 unpublished files from our lab. The details are listed in **Table S1**.

### Preparation of the training set for pre-filtering model

The 952 files were analyzed for precursor identification using DIA-NN (version 1.8.1), DIA-NN (version 1.7.12), and Spectronaut (version 14.6).

Target precursors identified by DIA-NN (version 1.8.1) and Spectronaut were used as positive samples, while decoy precursors identified by DIA-NN (version 1.7.12) served as negative samples. If the number of decoy precursors identified by DIA-NN (version 1.7.12) exceeded the total number of target precursors identified by DIA-NN (version 1.8.1) and Spectronaut, those decoy precursors with higher q-values were removed to balance the number of positive and negative samples.

Decoy precursors were acquired using DIA-NN (version 1.7.12), which can output decoy precursors along with the retention times (RTs)of their peak groups. DPHL v.2^18^ was utilized as the spectral library for library-based database searches. Decoy sequences were generated using mutation methods. The FDR threshold for all the database searches was set to 0.01.

For each precursor identified in each mass spectrometry file, peak groups were extracted based on the specified retention time (RT), as well as the precursor’s m/z and fragment m/z from DPHL v.2. Peak groups were extracted from MS1 spectra and MS2 spectra. Each peak was extracted over an m/z range of [ion m/z -20 ppm, ion m/z + 20 ppm] and covered 16 spectra (the spectrum corresponding to the precursor’s RT, plus seven spectra before and eight spectra after). The extracted peak group data from MS1 and MS2 were merged to create a 330×16 peak group matrix (the matrix was padded with zeros in the remaining rows if the precursor did not have enough fragments in the spectral library to fill the 330 rows). in addition to extracting peak groups centered around the spectrum indicated by the RT in the identification results, for each decoy, peak groups were extracted centered around a randomly selected spectrum within the [RT - 5, RT + 5] range as negative samples. This additional sampling helps the model classify diverse peak groups correctly during the pre-filtering process.

### Preparation of the training set for the pre-trained scoring model

Using the pre-filtering model, the best peak group for each precursor in the DPHL v.2 spectral library was determined in each mass spectrometry file. Specifically, a mapping function between iRT and RT was established for each mass spectrometry file. Then, for each precursor, the estimated RT was obtained based on its iRT in the spectral library using the mapping function. Peak groups were extracted for each spectrum within the [RT - n, RT + n] range and scored using the pre-filtering model. The peak group with the highest score for each precursor was retained. Precursors and their peak groups with scores above a certain experimentally determined threshold were kept as positive samples.

The peak group with the highest score for each decoy was selected. For each MS file, an equal number of decoys to the targets were chosen as negative samples.

### Preparation of the training set for the quantification model

A simulated dataset is prepared for the pre-training of quantification model. Empirically-corrected peptide library was build using EncyclopeDIA^25, 26^ (version 2.12.30) and Prosit^27^ based on uniport human fasta file (20,427 reviewed proteins, downloaded on 2023/10/26). For EncyclopeDIA, we used the convert tool panel to convert fasta to Prosit csv file by default parameters. For Prosit, we select “Prosit_2020_intensity_hcd” as intensity prediction model, and selected “Prosit_2019_irt” as iRT prediction model.

After obtaining the empirically-corrected peptide library, it was split into smaller libraries. The m/z was corrected using the method used in Synthedia (considering the mass of iodoacetamide modification and proton). Every four smaller libraries were randomly combined into a sub-library. For RT shift situations, the iRT of each fragment was randomly shifted negatively or positively. For peptide precursor permeation, we randomly selected a fragment ion and changed its m/z and charge to match with the peptide precursor’s m/z and charge for each peptide precursor in the smaller library. The peak group and label were extracted based on RT and TRUE area values recorded during simulation.

The quantification model was finally fine-tuned on a dataset extracted from 36 existing DIA-MS files of human cell line samples (PXD023139: WT and Ezh2-KO K562 cells; PXD033515: AML ex vivo cells). For each file, peak groups were extracted for target precursors identified by DIA-NN (version 1.9), and the sum of peak areas, serving as the label values, were extracted from the quantification result of DIA-NN (version 1.9). In our experiments, only the peaks of the first six fragments in the spectral library are included.

### Identification and quantification model

The identification model primarily consists of five components. Initially, for each row (ion) in the 330×16 peak group matrix, attributes from the spectral library are encoded into numerical features using numerical embeddings and concatenated with the peak group matrix and input to the model. Secondly, the input peak group matrix is processed through two convolutional layers, which include convolution, average pooling, and layer normalization. Thirdly, a transformer model with 8 self-attention blocks is applied to the matrix to capture long-range correlations, and then the matrix is downsampled to a 1D tensor through two fully connected layers. Fourthly, attributes of the current MS file are encoded into numerical features using numerical embeddings, and then downsampled into a 1D tensor through two fully connected layers. Finally, the above two 1D tensors are concatenated, and two fully connected layers are applied to map them to the sample label space through binary cross-entropy loss.

The architecture of our peak area estimation Transformer model mirrors that of the model used for peak group scoring, with the primary distinction being the use of a regression loss function to estimate the sum of peak areas. In our experiments, only the peaks of the first six fragments in the spectral library are included during training and identification.

### Precursor identification process

DIA-BERT employs a two-stage method for precursor identification in mass spectrometry files. The first stage is pre-filtering, which processes the mass spectrometry files similarly to the “Preparation of the training set for pre-trained scoring model” process, outputting a dataset composed of target and decoy precursors. In the second stage, the dataset is used to fine-tune a model initialized with the pre-trained scoring model. During fine-tuning, DIA-BERT splits the data into five parts to prevent overfitting by monitoring the decrease in validation set loss. DIA-BERT then re-scores the peak groups in the dataset using the fine-tuned scoring model and determines the identification results based on an FDR threshold of 0.01.

### Cross-run normalization

DIA-BERT enables cross-run precursor ion normalization. The intensities of precursors were firstly normalized by dividing the summed precursor ion intensity in each acquisition. Precursors were sorted by their indexed retention time (iRT) and grouped into bins of 100 precursors. For each bin, the median retention time (RT) across samples was calculated, and RT normalization was performed by subtracting these medians from the RT values. Then the RT-dependent normalization was conducted. For the initial sample, precursors were sorted by RT and grouped into bins of 400. For subsequent samples, precursors were assigned to bins based on the RT boundaries defined by the first sample. The difference in bin rank between samples was calculated, and quantification values were marked as missing (NA) if the bin-difference exceeded 2. The bin ranks for each precursor across samples were adjusted to the median rank. Within each RT bin, precursor quantification was normalized by dividing by the median quantification value of that bin in each sample. Protein quantification was then performed using MaxLFQ ^28^ utilizing the RT-dependent normalized precursors. The resulting protein quantification was used to further adjust precursor quantification. For each precursor within a protein, the intensities across samples were summed, and the summed values were converted to percentages, which served as adjustment factors. The adjusted precursor quantification was obtained by multiplying the protein quantification by these factors. Finally, any precursor quantification associated with NA values in the RT-dependent normalized matrix was also set to NA.

### Benchmark experiment

For identification comparisons, including the identification of peptide precursors and proteins, as well as the quantification correlations, each benchmarking file was independently searched using DIA-NN (version 1.9). For quantification comparisons, proteome files from three species (n = 6) were processed with DIA-NN (version 1.9) using combined search with MaxLFQ normalization. The parameters were set as ‘--qvalue 0.01 --cut K*,R* --min-fr-mz 200 --max-fr-mz 1800 --missed-cleavages 1 --min-pep-len 7 --max-pep-len 30 --min-pr-mz 300 --max-pr-mz 1800 --min-pr-charge 2 --max-pr-charge 4 --unimod4 --var-mods 1 --var-mod UniMod:35,15.994915,M --relaxed-prot-inf --rt-profiling --pg-level 0’. For the library-based database search, the DPHL v.2 spectral library was used for the human proteome search, while a spectral library from our previous work^19^ was used for the three-species database search.

## Acknowledgments

This study and the coauthors of this study are jointly supported by grants from the National Key R&D Program of China National Key R&D Program of China (No. 2021YFA1301600), and the Westlake Educational Foundation. We thank the Pioneer and Leading Goose R&D Program of Zhejiang (2024SSYS0035). We thank Westlake University Supercomputer Center for assistance in data analysis and storage.

## Conflict of interests

T.G. is the founder of Westlake Omics (Hangzhou) Biotechnology Co., Ltd., while P.L., and Z.N. are staff of this company.

## Availability the data and codes

Codes and data relevant for data analysis in this study is available at https://github.com/guomics-lab/DIA-BERT. The executable version of DIA-BERT and manual are available at http://dia-bert.guomics.com/index.html.

## Author contributions

T.G. acquired funding for and supervised the project. T.G. and Y.C. conceptualized the research and provided supervision. Y.C. designed the methodology and setting the stage for the research implementation. P.L. implemented the overall experiment pipeline. Z.N. developed the data preprocessing component and DIA-BERT software system. Z.L. implemented the simulation component and produced the simulated data. Z.L., Z.N., and Y.S. curated the data. P.L., Z.L., Z.N., Y.S., X.Z. and Y.Z. conducted the model parameter tuning and carried out analyses. Z.L. devised benchmark experiments and drafted initial manuscript with contribution from P.L. and Z.N.. Z.L., T.G., Y.C., and Y.S. revised the manuscript. All authors have read and approved the final manuscript.

## Supplementary Tables

Table S1: Training files and comparison summary of identification between DIA-NN and DIA-BERT Table S2: Identified peptide precursors and proteins using DIA-NN (in library-based mode) and

DIA-BERT

Table S3: Parameters for simulated data by modified Synthedia

**Figure S1.**
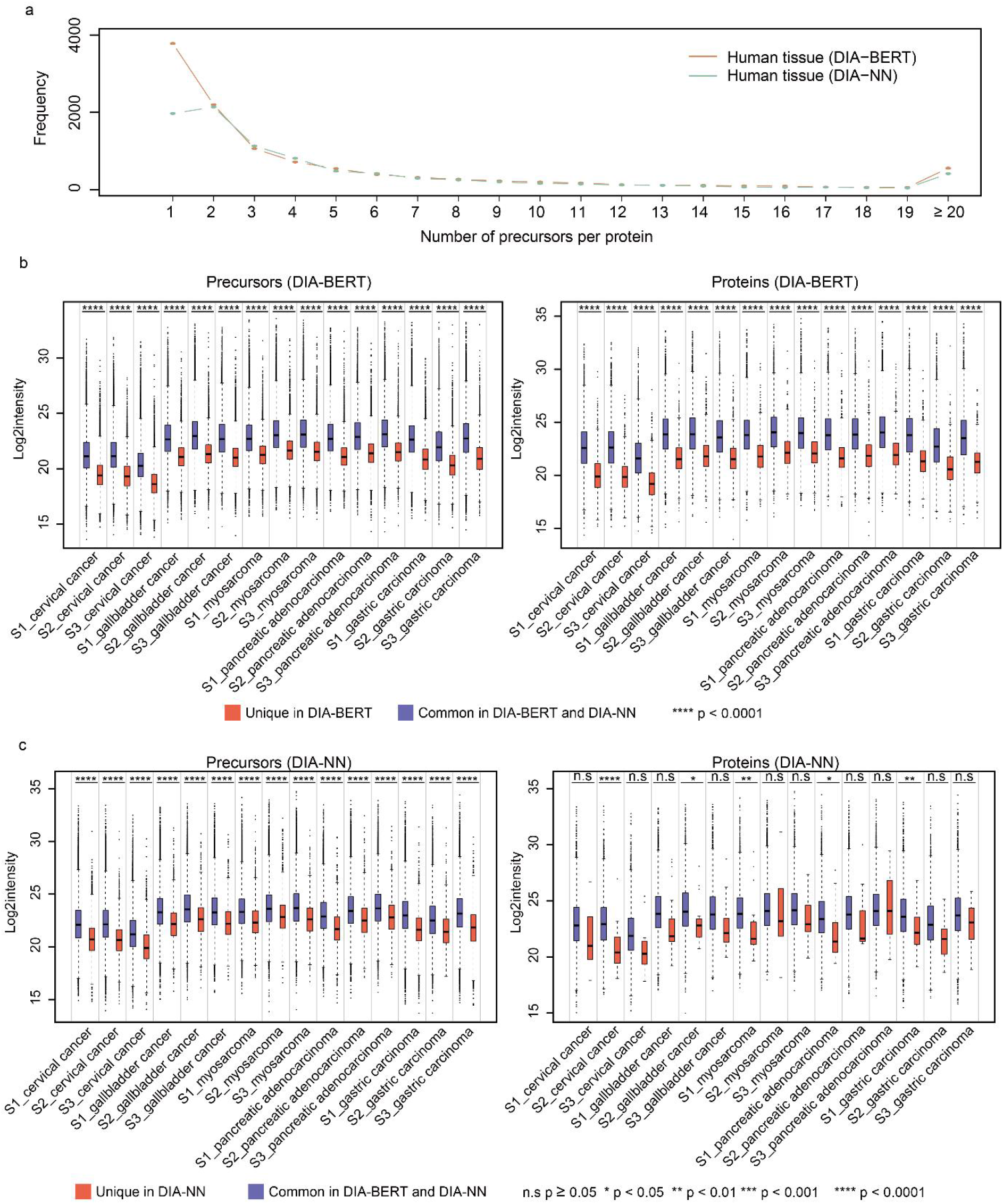
Comparison of peptide and protein identifications in human proteome data set. **a**, The frequency of the number of peptide precursors per protein in DIA-BERT and DIA-NN. **b**, The significant abundance difference between unique peptide precursors/proteins identified by DIA-BERT and common peptide precursors/proteins identified by both of DIA-BERT and DIA-NN. Whiskers, 1.5x interquartile range. **c**, The significant abundance difference between unique peptide precursors/proteins identified by DIA-NN and common peptide precursors/proteins identified by both of DIA-BERT and DIA-NN. Whiskers, 1.5x interquartile range.

**Figure S2.**
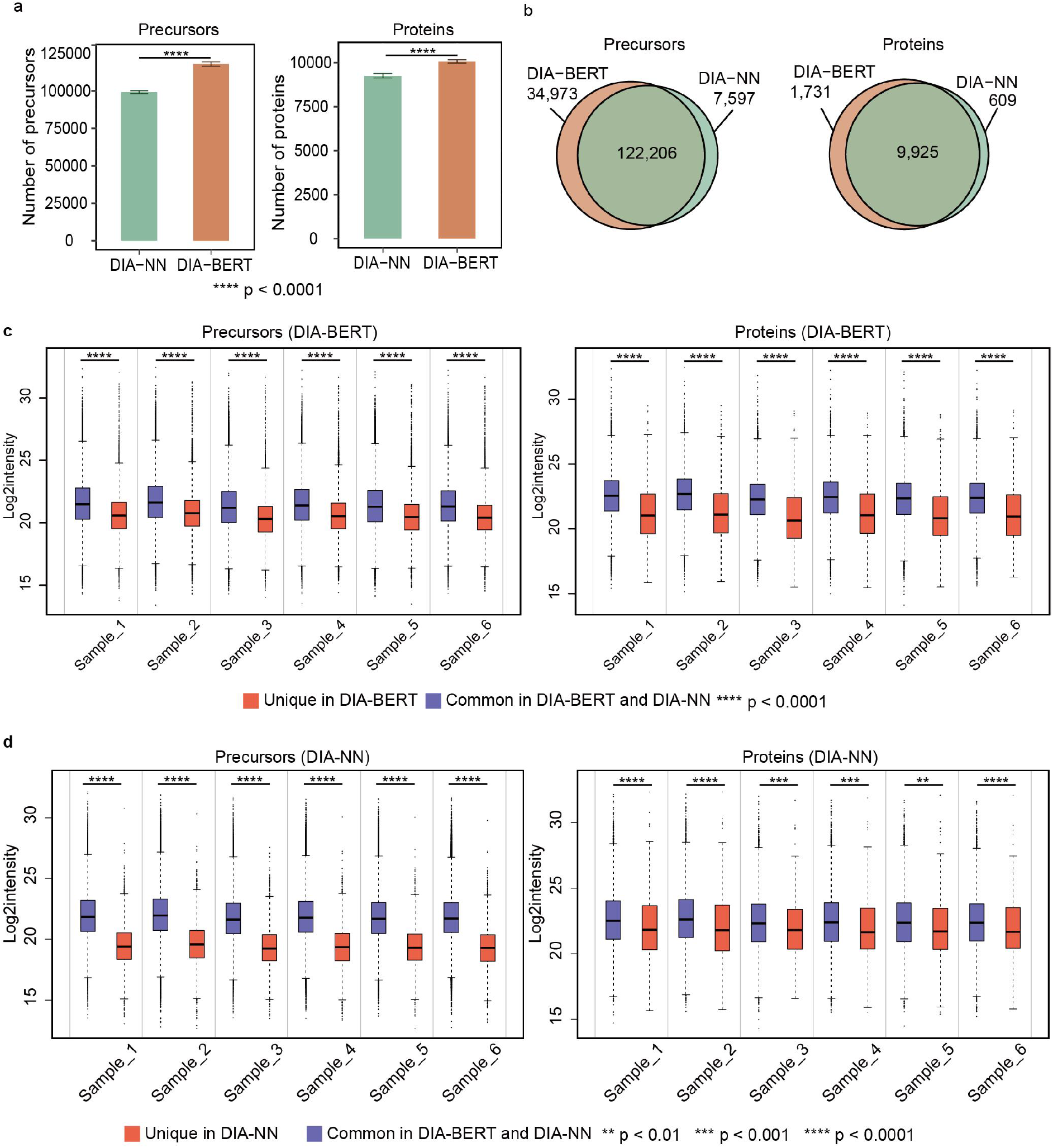
The identification comparison in three species proteome data set. a, The identification comparison between DIA-BERT and DIA-NN in three species proteome data set for each sample. **b**, The overlap of identified three species peptide precursors or proteins using different software. **c**, The significant abundance difference between unique peptide precursors/proteins identified by DIA-BERT and common peptide precursors/proteins identified by both of DIA-BERT and DIA-NN for three species proteome data set. Whiskers, 1.5x interquartile range. **d**, The significant abundance difference between unique peptide precursors/proteins identified by DIA-NN and common peptide precursors/proteins identified by both of DIA-BERT and DIA-NN for three species proteome data set. Whiskers, 1.5x interquartile range.

**Figure S3.**
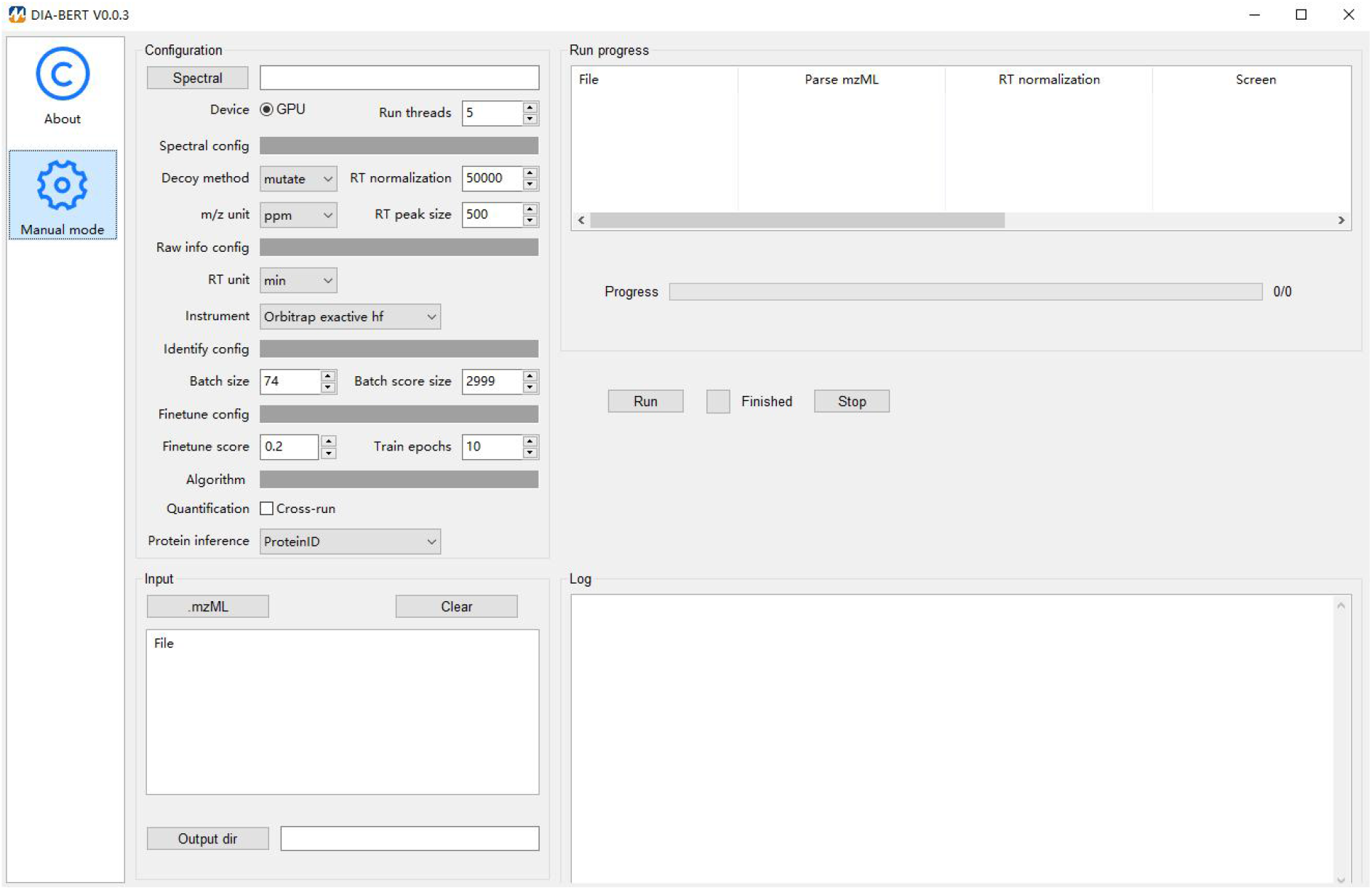
The GUI of DIA-BERT. The Input section allows users to specify paths for raw files (mzML format) and customize the output directory (.csv format). The Configuration section enables adjustments for settings such as the spectral library, run threads, decoy method, RT normalization, m/z unit, RT peak size, RT unit, instrument, batch size, batch score size, RT width, fine-tune score, cross-run quantification, and the number of epochs for model optimization. The Run Progress section monitors the analysis, allowing users to toggle between run and pause states while providing real-time progress updates. The Log section serves as a comprehensive runtime log, capturing essential events, state transitions, errors, warnings, and critical runtime details.

